# From zygote to a multicellular soma: body size affects optimal growth strategies under cancer risk

**DOI:** 10.1101/2019.12.16.877118

**Authors:** E. Yagmur Erten, Hanna Kokko

## Abstract

Multicellularity evolved independently in multiple lineages, yielding organisms with a wide range of adult sizes. Building an intact soma is not a trivial task, when dividing cells accumulate damage. Here, we study ‘ontogenetic management strategies’, i.e. rules of dividing, differentiating and killing somatic cells, to examine two questions. First, do these rules evolve differently for organisms differing in the target mature body size, and second, how well a strategy evolved in small-bodied organisms performs if implemented in a large body – and vice versa (‘large’-evolved strategies in small bodies). We model the growth and mature lifespan of an organism starting from a single cell and optimize, using a genetic algorithm, trait combinations across a range of target sizes, with seven evolving traits: 1) probability of asymmetric division, 2) probability of differentiation (per symmetric cell division), 3) Hayflick limit, 4) damage response threshold, 5) damage response strength, 6) number of differentiation steps, 7) division propensity of cells relative to ‘stemness’. Some, but not all traits, evolve differently depending on body size: large-bodied organisms perform best with a smaller probability of differentiation, a larger number of differentiation steps on the way to form a tissue, and a higher threshold of cellular damage to trigger cell death, than small organisms. Strategies evolved in large organisms are more robust: they maintain high performance across a wide range of body sizes, while those that evolved in smaller organisms fail when attempting to create a large body. This highlights an asymmetry: under various risks of developmental failure and cancer, it is easier for a lineage to become miniaturized (should selection otherwise favour this) than to increase in size.

## Introduction

Multicellularity allows an organism to divide tasks between cells, creating a complex soma with a much wider range of functions than single-celled organisms can achieve (Grosberg and Strathmann, 2007; Knoll, 2011). It also permits large body size, which may help in predator avoidance (Boraas et al., 1998; Kapsetaki and West, 2019) and be subject to other types of selection to become large, including metabolic scaling (White et al., 2019) and intraspecific competition (Kingsolver and Pfennig, 2004; Pettersen et al., 2015). However, creating and maintaining the soma of a multicellular organism is not a trivial task. This requires cooperation between their individual cells, where somatic cells forgo passing their genetic material to the offspring, and they divide only to maintain organismal functions. From a single somatic cell’s perspective, if the benefits of cooperating outweigh self-replicating, this can still be advantageous (Goldsby et al., 2014; Libby et al., 2016). But even if somatic cells ‘aim’ to cooperate, they inherit their genomes from germline cells that have unlimited dividing potential, and the adjustments required to make cells a differentiated part of a somatic tissue may not always function flawlessly. Uncontrolled growth (cancer) is one problem (Aktipis et al., 2015), and since tumours typically only arise when several mutations have occurred in the same cell lineage, the number of ‘permitted’ cell divisions has evolved to be limited (Sager, 1991; Campisi, 2013; Maciejowski and de Lange, 2017). Together with other aspects of somatic maintenance, such as damaged cells being recognized and entering cell cycle arrest or undergoing programmed cell death (Fuchs and Steller, 2011; Campisi, 2013), and the hierarchical tissue organization (Derényi and Szöllősi, 2017) that maintains some cells in a stemlike state while differentiates others, these rules reflect what we call the ‘ontogenetic management strategy’ of an organism’s cells.

There are good reasons to suspect that such a strategy is not of a ‘one size fits all’ type, i.e., that ontogenetic rules evolve to be different depending on the target body size at maturity. Despite the great range of body sizes in nature, many multicellular organisms go through a single-cell stage. The resulting high relatedness between cells appears to be a way to improve cooperation between individual cells (Queller, 2000; Grosberg and Strathmann, 2007; Fisher et al., 2013; Bastiaans et al., 2016), but it also means that larger organisms need to do more cell divisions to reach their mature size. Moreover, once mature, the adult size needs to be maintained, balancing the cell turnover that occurs throughout life (Fuchs and Steller, 2011; Galluzzi et al., 2018); again, larger organisms maintain a larger cell population. Mature body size is also not the only axis of variation that is likely to matter for the optimal strategy: extrinsic mortality can also matter (Boddy et al., 2015). Organisms that face a high risk of extrinsic mortality need to ensure that they mature fast enough to reproduce before they die. Low-mortality scenarios in turn give prolonged opportunities to reproduce (accumulating fitness iteroparously), but these can only be realized by organisms with robust enough bodies that last a sufficiently long time.

Empirical evidence for variation in traits that comprise the ontogenetic strategy can be found in both comparative ageing and oncology literature. Already in the mid-20th century, Hayflick and Moorhead observed that somatic cells could only divide a finite number of times, termed the ‘Hayflick limit’ (Hayflick and Moorhead, 1961; Hayflick, 1965; Shay and Wright, 2019). This limit is related to telomere length and telomerase expression, and both affect cancer-and senescence-related processes, with e.g. telomerase expression observed in most human tumours (Kim et al., 1994), allowing tumour cells to divide without hitting the Hayflick limit (Blasco, 2005; Shay and Wright, 2019). Comparative studies have revealed correlations between body size and/or lifespan and telomerase activity and telomere length (Seluanov et al., 2007; Gomes et al., 2011), a pattern also predicted by theory (Risques and Promislow, 2018). For example, the finding that telomerase expression is repressed in somatic tissues of humans and larger mammals (Kim et al., 1994; Gomes et al., 2011) fits in well with the idea that telomerase expression is not a risk-free way to increase lifespan. It enables cell lineages to persist for longer, but may open the door to cancer. Again, this may be a problem for large organisms in particular, as they need many cells for a long time, while careless attempts to do this, e.g. via increasing the Hayflick limit, may end their lives with cancer instead.

The finding that cancer incidence does not scale with body size despite an increased number of cell divisions (Peto’s paradox: Peto et al., 1975) has led to a quest to find the adaptations that mitigate cancer risk as body size increases (Caulin and Maley, 2011). In addition to telomere management, potential adaptations include stronger DNA damage response (for a comparison of elephants and humans see Abegglen et al., 2015), or increasing the number of differentiation levels from the stem cell to the terminally-differentiated cells (theory: Derényi and Szöllősi, 2017). The precise manner in which cells populate these degrees of differentiation has implications for controlling damage accumulation in tissues and consequent cancer risk (e.g. Shahriyari and Komarova, 2013), as well as tissue homeostasis and senescence-related processes (Liu et al., 2019).

Our aim is to combine the above considerations in a single model. The various mechanisms listed above do not act in isolation, instead, they orchestrate ontogeny and survival in concert with each other. We model organismal growth and subsequent mature lifespan, assuming throughout that the organism starts its life as a single cell and reaches its target mature body size through divisions and differentiations (unless it dies beforehand through extrinsic mortality or ‘mismanagement’ of its own growth). We consider the trade-offs inherent in any ontogenetic strategy: delaying death via one cause (e.g., by increasing the Hayflick limit) may hasten the probabilistic death from other causes (e.g., increased cancer risk). The interacting causes of death allow us to quantify the expected fitness of any given ontogenetic management strategy. We specifically investigate whether a genetic algorithm finds the same solutions across different body sizes, or larger bodies require fundamentally different ontogenetic management than small bodies. Some traits turn out to be more body size specific than others, such that as a whole, multiple alternative combinations of traits may yield a well-functioning soma for a given body size. Finally, we investigate how the cell-management strategies that evolved with specific body sizes fare across a wider range of body sizes. Here, a clear asymmetry emerges: ontogenetic management strategies that evolved in smaller organisms generally fail to perform in large organisms, leading to frequent premature death. The converse is not true: strategies adapted to a large body size fare reasonably well across all body sizes.

## Materials and Methods

### Model overview

Our model tracks the number and state space of all the cells of an organism (see Fig.1 for an overview) from zygote to organism-level death. Deaths occur stochastically, thus the success of a strategy is evaluated by running independent realizations of the growth and lifespan that results from a specific ontogenetic strategy. The ontogenetic strategy has a total of seven components, all evolving within a prespecified plausible range (see Table 1 for permitted ranges of values): the Hayflick limit (the number of divisions a cell can go through), denoted *H*; the number of levels to terminal differentiation, *T*; the probability that a cell division is asymmetric, *P*; the differentiation probability in symmetric divisions, *Q*; the division propensity of each differentiation level (relative to the previous level), *X*; the DNA damage response threshold, *A*; and the probability that a cell is killed upon DNA damage detection, *S*.

**Figure 1:**
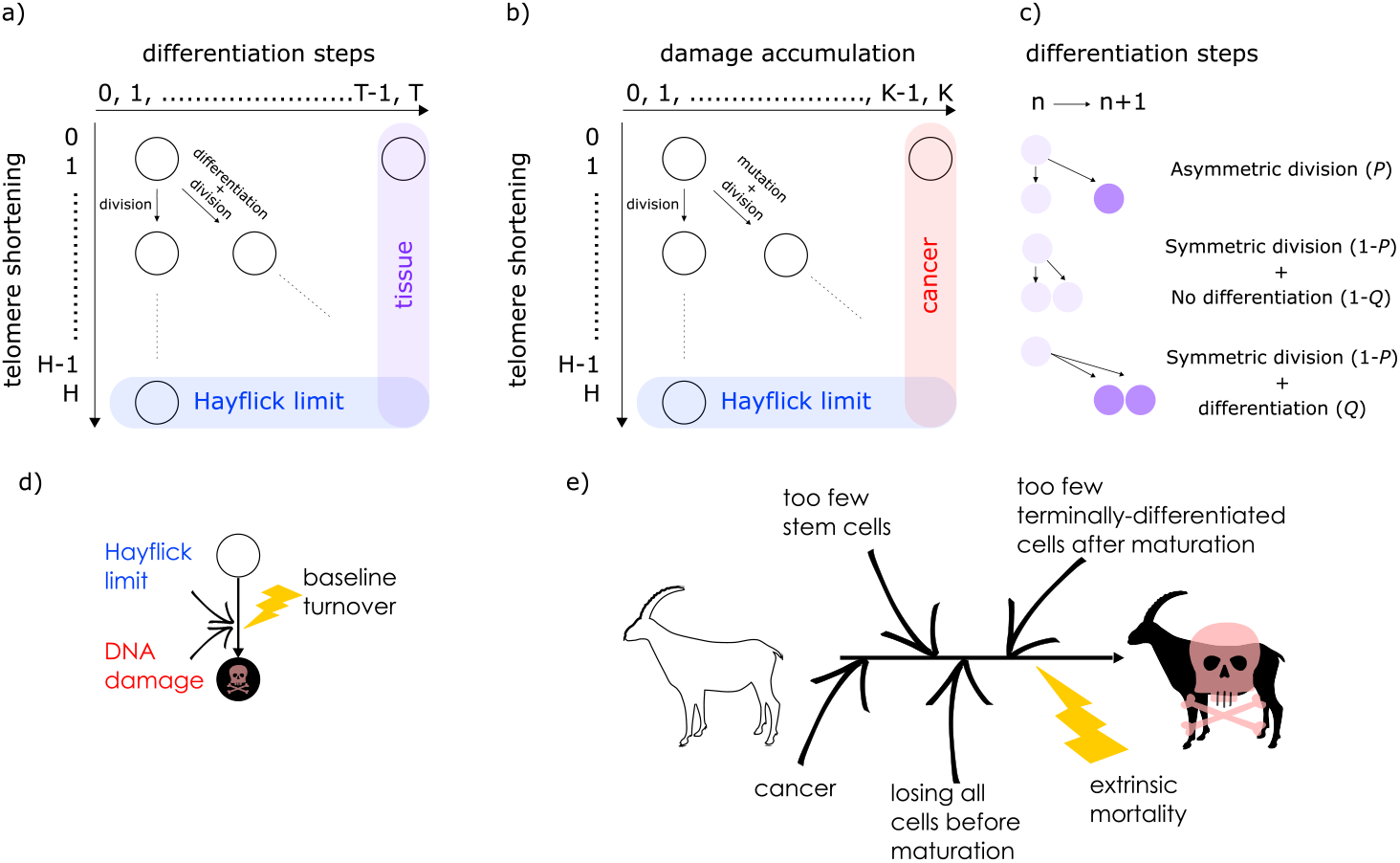
The model structure. In panels (a) and (b), rows represent the number of divisions a cell lineage has accumulated (since the zygote). As cells divide, their telomeres shorten, which moves them down to the next row. Any cell that reaches the last row is removed (loss of telomeres; blue in both panels). The two other states are (a) differentiation towards tissue (reached at terminal differentiation, *T*) and (b) mutations causing DNA damage. Unlike the shortening of telomeres, these two aspects happen probabilistically, with the organism controlling differentiation via its ontogenetic strategy (c), and mutations occurring with probability *c* in the new cells. A cell that has reached the final column (purple) in (a) cannot divide further but contributes to somatic function; a cell that reaches the final column (red) in (b) becomes oncogenic and kills the organism. (d) Cellular turnover occurs as a result of various processes of cell removal: telomere loss, DNA damage-related cell death, or a baseline turnover rate *ν*. (e) The organism can die due to various causes related to extrinsic mortality, failed ontogenetic management of its cell population, or cancer.

**Table 1:** Traits that form the ontogenetic strategy, together with the possible range within which evolution can take place (new mutations occur within the prespecified range)

| Symbol | Trait | Range |
| --- | --- | --- |
| $P$ | Probability that a cell division is asymmetric | real numbers between 0.0 and 1.0 |
| $Q$ | Differentiation probability in symmetric divisions | real numbers between 0.0 and 1.0 |
| $T$ | Number of differentiation steps | integers between 3 and 50 |
| $X$ | Division propensity of each differentiation level (relative to the previous level) | real numbers between 0.01 and 200 |
| $S$ | Percent of cells killed upon damage detection | real numbers between 0.0 and 1.0 |
| $A$ | Damage response threshold | real numbers between 0 and $K^*$ |
| $H$ | Hayflick limit (the maximum permitted number of cell divisions) | integers between 5 and 500 |

\* $K$ is the number of oncogenic steps that, if completed, result in cancer

The individual organism is modelled as a population of cells. At each time step, the organism experiences a cell number deficit *D* if it has fewer terminally differentiated cells than its body size target (details on how *D* is determined are given below). Filling this deficit requires cells to divide, and this can change the state of the participating cells in three different ways. We do not give spatial coordinates or specific tissue functions to cells, instead cells may differ from each other in three distinct (but possibly covarying) ways: how differentiated they are, how many divisions they are a result of (since the zygote state), and how damaged they are. The variables *H* to *S*, above, interact to determine the transition probabilities between these states. We give a brief description of the three-dimensional state here, and describe the actual interactions between variables and movements in state space later.

First, cells differ in their level of differentiation (horizontal axis in Fig.1a), modelled as integer-valued levels from 0, the zygote, to *T*, a cell that contributes to tissue. All non-terminally differentiated cells are hereafter collectively called ‘stemlike cells’ (with total number *N*_stemlike_), which include the actual stem cells (level 0, *N*_0_) as well as the more differentiated transit-amplifying cells, *N*_TAC_, that still have division potential (levels 1 to *T*–1). Terminally differentiated cells (level *T*, *N*_tissue_) perform somatic functions and cannot be chosen to divide further. The value of *T* evolves, i.e. is part of the ontogenetic strategy.

The second component of a cell’s state is the number of times the cell has divided from the zygote state (shown vertically in both Fig.1a and 1b; note that the entire cell state structure is three-dimensional). Values range from 0 (the zygote) to *H* (the Hayflick limit: Hayflick and Moorhead, 1961; Hayflick, 1965). Although we do not track declining telomere length explicitly (in terms of base pairs), a cell moving along this axis can be interpreted as losing telomere length, such that cells that have reached level *H* have too short telomeres to divide further. These cells are removed whether or not they are also terminally differentiated. The value of *H* itself corresponds to telomere length at the beginning of life (specifying the maximum number of cell divisions permitted for any cell lineage thereafter), and is part of the ontogenetic strategy.

The third component of cell state (horizontal axis in Fig.1b) tracks the accumulated damage in a cell, corresponding to oncogenic steps termed ‘mutational hits’ or ‘rate-limiting steps’ in models (Nunney, 1999; Calabrese and Shibata, 2010; Kokko and Hochberg, 2015). We track damage with integers from 0 to *K*, with 0 being the state without any oncogenic damage. The maximum tolerable damage, *K*, is not an evolving part of the ontogenetic strategy, but a constraint that organisms must evolve to deal with: once an organism has at least one cell of damage level *K*, it is assumed to die from cancer.

Cancer, however, is not the only way for an organism to die (Fig.1e). Life can end with 1) not having stemlike cells to keep dividing to maintain its differentiated cells, 2) falling below a certain number of terminally-differentiated cells after maturation, 3) losing all cells before maturation, 4) cancer, and 5) extrinsic mortality (parameter *μ*) (Table 2). We further divide the death cause 1 into pre-and post maturation, to distinguish between developmental (1-a) and somatic maintenance failures (1-b). Some of the causes of death in this list, such as failure of the developmental program to produce any terminally-differentiated cells, are less likely than others to be observed in real organisms. However, since our aim is to investigate which ontogenetic strategies work (i.e. can produce an organism that can mature and reproduce) and which do not, we need to explicitly track deaths due to failures to achieve a functioning soma in the first place.

**Table 2:** Death causes for an individual organism

| Cause | Criterion | Number in the text |
| --- | --- | --- |
| Depleting stemlike cells pre- or post-maturation | $N_{\text{stemlike}} = 0$ | 1-a & 1-b |
| Not having enough tissue cells (after maturity) | $N_{\text{tissue}} < M \times 0.8$ | 2 |
| Losing all the cells before reaching maturation | $N = 0$ | 3 |
| Cancer | at least one cell<br>at damage level $K$ | 4 |
| Extrinsic mortality ($\mu$) | $t \geq -\ln 0.001/\mu$ | 5 |

A zygote begins its life in the state 0, 0, 0 (no differentiation, no divisions yet, no damage accumulation). We track the subsequent number of cells in the entire 3-dimensional state space over time (the age of the organism) as the fourth dimension. At death, we record the fitness of the organism. No fitness accumulates before reaching mature size (a parameter, *M*, that we vary over 2 orders of magnitude); afterwards, each time step contributes to cumulative fitness. Since only terminally differentiated cells are assumed able to perform useful functions for the whole organism, we model the per-step fitness gain as the proportion of terminally differentiated cells (out of all cells) that the organism currently has. This reflects the biologically plausible assumption that cells that are not terminally differentiated impose maintenance costs while not yet contributing to somatic function. Maintaining an excessive population of such cells therefore reduces fitness. The expected fitness of the ontogenetic strategy (as opposed to a single realization, i.e. an individual life) is obtained by rerunning this process multiple times.

### Growth, differentiation, and fitness

Here we describe the procedures that create a single run of an organism’s life. Starting from a single cell (zygote), the organism grows via division and differentiation of its cells. It matures when it has reached a predefined size called the maturity threshold *M* defined as a sufficient number of terminally differentiated cells. We track an individual’s fitness accumulation until *t*_death_ where continuing life is not possible for death reasons 1 to 4 listed in the Table 2, or when it has reached an age *t*_end_ = −ln0.001/*μ*, an age that is only reached with probability 0.001 when extrinsic mortality (the fifth cause of death) equals *μ*. If the organism reached maturity before dying, its age at this point is recorded as *t*_maturation_. Thereafter, the expected fitness is computed as a weighted sum of the fitness gain.

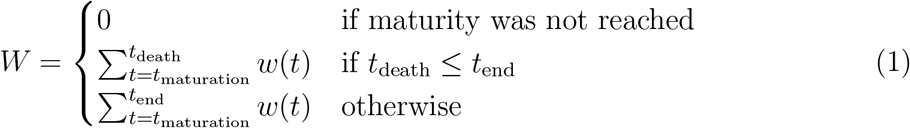

Here 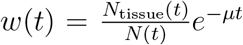, where *N*_tissue_(*t*) gives the numbers of terminally differentiated cells at age *t*, *N*(*t*) is the total number of cells (= *N*_tissue_(*t*) + *N*_stemlike_(*t*)), and the last term is the probability that the organism has escaped extrinsic mortality up to age *t*.

To find *w*(*t*) and the times of maturity and death, we need to describe how the ontogenetic strategy spreads new cells into the state space. We assume that an organism grows towards the maturity threshold before it has reached this size, and aims to maintain the number of terminally differentiated cells thereafter (which requires compensating for cells that have died either because of a baseline cell turnover *ν*, see below, or because of being killed as part of the ontogenetic strategy). Thus, both before maturity and afterwards, the organism begins a time step by measuring the difference between the target number of terminally differentiated cells (*M*) and its current number (*N*_tissue_). The deficit *D* is then defined as *D* = *k*(*M* – *N*_tissue_), rounded up to the nearest integer, where the multiplication with a constant *k* makes growth before maturity approximate a von Bertalanffy trajectory (Karkach, 2006).

In case of a positive deficit, up to *D* stemlike cells will divide. If *D* ≤ *N*_stemlike_, the number of dividing cells is *D*, which makes the deficit vanish. If *D* > *N*_stemlike_, then all *N*_stemlike_ cells divide, and the deficit diminishes. In both cases, *D* will be calculated anew in the next time step. If there is an excess of stemlike cells to choose from (the first case), the precise identity of cells chosen for division follows component X of the organism’s ontogenetic strategy, in the following manner.

Stemlike cells occur in different states of differentiation 0, 1,…, *T*–1. Ontogenetic strategy component *X* is defined as the relative division propensity of a given level of differentiation compared to the previous one (*X* > 1 means that propensities increase with level of differentiation). Cells are chosen (without replacement) to divide with relative propensities that are proportional to *X^i^* where *i* is the cell’s current degree of differentiation, until the appropriate (see above) number of cells have been chosen. A cell’s position along the *H* or *K* axis does not directly impact its likelihood of becoming a dividing cell, though patterns of covariation are likely to arise because cells far on their path to full differentiation are also likely to have progressed further from the 0 state along these axes too. Thus if *X* is high, divisions use stem cells sparingly, which also means that divisions occur more commonly in cells that have already incurred some damage.

Once the state-specific numbers of dividing cells is known, each division is performed taking into account strategy components *P* and *Q* (Fig.1c). The division occurs asymmetrically with probability *P*, in which case one daughter cell is placed into the soma one step further along the differentiation axis while the other daughter cell does not change in its level of differentiation (for placements along the other axes, see below). With probability 1 –*P*, the cell divides symmetrically, and in this case component *Q* of the strategy is also relevant. Each of the two daughter cells enters the soma as either one step more differentiated than the parent (with probability *Q*, applied independently for both daughters) or at the same level of differentiation as the parent (with probability 1–*Q*, interpretable as replenishment of the pool of stemlike cells). Thus, as a whole, with probability (1 – *P*)*Q* a division occurs symmetrically and yields in two daughter cells that are both more differentiated than their parent, whereas with probability (1 – *P*)(1–*Q*) it yields in two cells at the identical level of differentiation as the mother cell, and finally with probability *P* the division produces one daughter that is more differentiated than the parent and another daughter that remains at the original differentiation level.

We also need to specify the positions of the new daughter cells along the *H* and *K* axes. Since the *H* axis tracks the number of cell divisions, the new location along this axis is always one step further than the old location (and since only stemlike cells, up to *H*-1, can be chosen to divide). Regarding DNA damage, each new daughter cell can be at the same location along the *K* axis (with probability 1–*c*) or with an increased level of DNA damage, i.e. one step further along the *K* axis (probability *c*).

At the end of a division round, we apply various sources of mortality to cells (Fig.1d). Parental cells have at this point become two daughter cells, necessitating removing the original parent from the soma. Cells that have reached the Hayflick limit (state *H*) are also removed. Furthermore, the ontogenetic strategy has two components, *A* and *S*, that aim to remove damaged cells (cells that are far along the *K* axis regardless of their position along other axes), but we assume that the recognition of damage is imperfect. A proportion *S* of cells with *a* + ϵ ≥ *A* are killed, where *a* is a cell’s true location along the *K* axis and an error term ϵ ~ *N*(0, 0.5) reflects the difficulty of determining precisely how damaged a cell is. Since recognized damage is a real number (not restricted to integer values), we also allow A to take real number values. The organism cannot evolve a smaller ϵ, but it can evolve a suitable *A*, with a clear trade-off: a low *A* is good for reliable removal of all damage, but a large number of healthy cells will then be removed too, depleting the pool of stemlike cells potentially too quickly. Finally, any of the cells that survive the above mortality risks die with probability *ν*, reflecting baseline cellular mortality (cell turnover), applied independently to each cell.

### Finding the optimal strategies using a genetic algorithm

The multidimensionality of the fitness landscape (seven evolving traits) creates a challenge for any optimization procedure, and we thus implement a genetic algorithm to seek for the best combinations of traits. We begin the optimization procedure with a population of 100 ancestral ontogenetic strategies, initializing each of the seven components (*P, Q, T, X, S, A* & *H*, Table 1) by picking from uniformly distributed ranges (the relevant ranges for each variable are listed in Table 1). Then, we estimate the fitness of each strategy (*W*, eq. 1) 10 times, but to speed up the optimization, we stop evaluating the fitness of a strategy if the first four estimates are zero, but nevertheless keep it in the population (with fitness estimated as zero) for the subsequent genetic algorithm run. For all other strategies, we estimate mean fitness 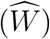 as the average value of W across the 10 runs. This set forms the initial population of candidate strategies.

We next start the genetic algorithm, consisting of numerous rounds. In each round we first rank the strategies with respect to their 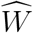 and pick 10 as ‘potential parents’ (in the sense of a genetic algorithm). To avoid the danger of populations becoming stuck on a local fitness peak, which results easily if the algorithm is allowed to focus on the 10 best strategies that each recombine with each other and thus become similar, we choose 10 equally spaced strategies from the population (i.e. 1st, 12th, 23rd,…,100th). We then proceed to form two ‘parents’ out of the potential parents. The probability that a strategy is selected as the first parent is proportional to its 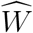. Once the identity of the first parent is known, we mutate it as follows: we create a novel random strategy within bounds shown in Table 1 and calculate the difference in trait values between the novel strategy and the parent strategy. The parent’s trait values are then shifted somewhat toward the new random strategy: for each component of the strategy, the new value is *x*_new_ = (1-*s*)*x*_old_ + *sx*_novel_, where *s* ~ Gamma(0.25, 0.1) truncated between 0 and 1. The parameters of the gamma function (shape=0.25, scale=0.1) are chosen to make sure that mutations generally happen with small steps, with occasional bigger jumps. Traits *T* and *H* are thereafter rounded to the nearest integer.

Genetic algorithms typically use a stylized version of crossovers of chromosomes. We use crossover by assuming that each ontogenetic strategy is represented with a string (formed by traits *P-Q-H-A-S-T-X*) analogous to a chromosome. We pick a random mid-point (between any two traits) and recombine the chromosomes of two parents, by picking a second parental strategy from the remaining pool of potential parents (this parent is chosen randomly, i.e. without taking into account relative fitness, and also is not mutated). In the resulting ‘offspring’ strategy, all the values up to that mid-point comes from one of the parents (either the newly mutated one, or the second parent, with 50% probability for either option), and the rest from the other parent.

The offspring strategy’s fitness 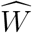 is evaluated using the same procedure as was done for the parental generation, except now we drop this offspring (and go back to creating a new offspring with the procedure above) if at any point after the first four of the ten realizations the mean of its *W* values fall below the parental population’s best 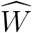 (as such an offspring is unlikely to be a winner). If the offspring’s 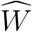 based on all 10 realizations exceeds the entire population’s best-ranked strategy’s 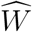, it becomes the new ‘best strategy’ and the lowest-performing strategy is removed from the population. In the alternative case where the offspring’s 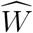 is lower than the best, but still better than the worst-performing 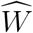 present in the population, we add the offspring strategy to the population as a potential parent that could be picked in the next rounds (and again remove the worst one).

We stop the genetic algorithm if no new ‘best strategy’ is found for 300 rounds. We run the genetic algorithm four times independently for each maturity threshold *M*, from different initialization points of the original population.

## Results

Optimized life histories reached much higher lifetime fitness at small than at large body sizes (Fig.2a). While surprising at first sight, note that our model did not give large organisms any of the advantages often associated with body size (Kingsolver and Pfennig, 2004). We therefore do not examine selection to become large per se, but ask how organisms deal with the increased difficulties that creating a larger body entails. A major challenge is that reaching maturation size takes more time if the required size is larger (Fig.2b). Meeting this challenge requires changing some, but not all, components of the ontogenetic strategy in a systematic manner. The probability of differentiation (*Q*) declines with body size (Fig.2d), while the number of levels to terminal differentiation (*T*) increases (Fig.2e). Also, at larger body sizes, the damage response threshold (*A*) was never found to be low, whereas low body sizes can be viable with a range of values for A (Fig.2h).

**Figure 2:**
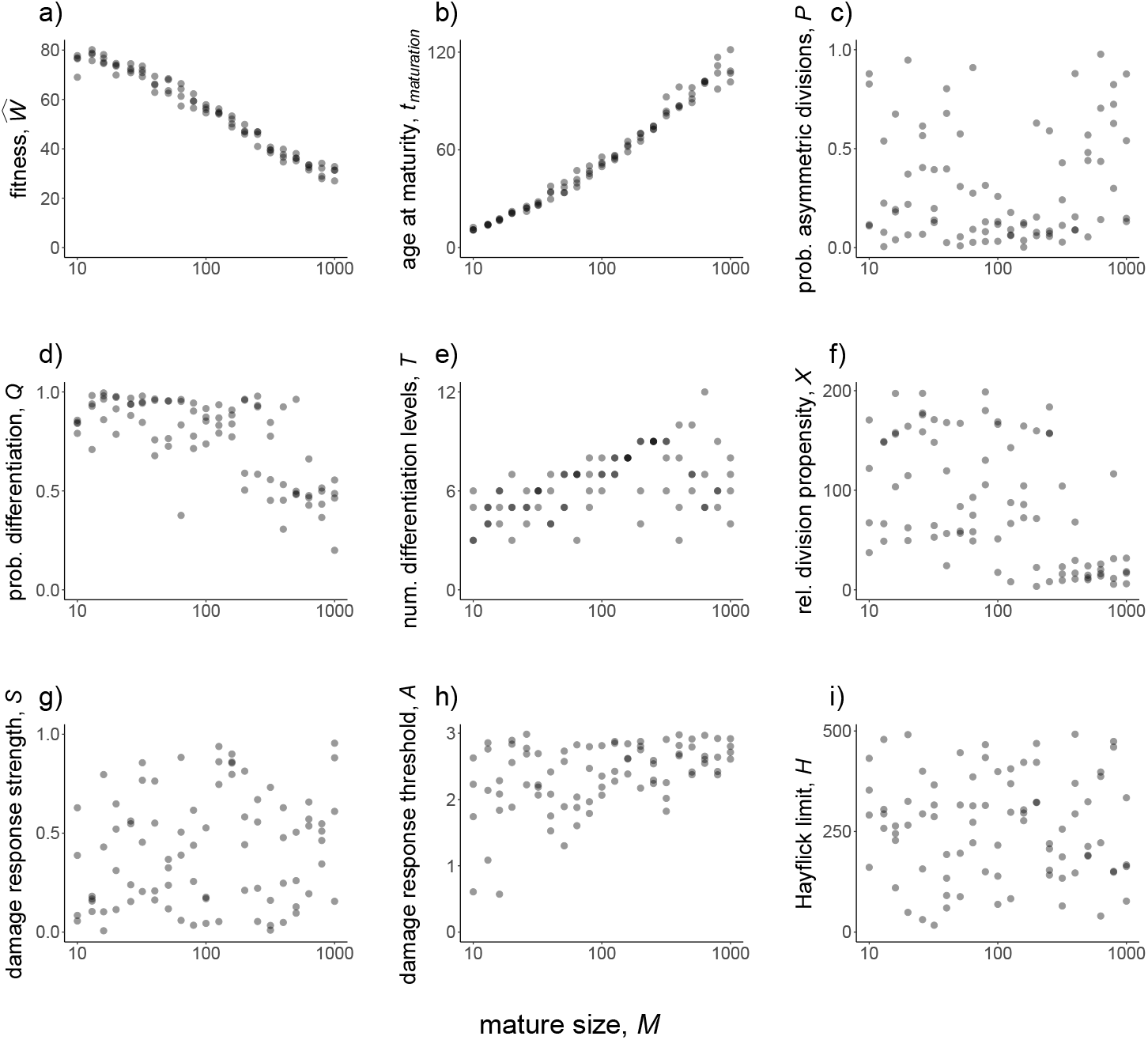
Properties of optimized ontogenetic strategies across 21 logarithmically spaced body sizes. Data refer to the best strategy at the end of each optimization procedure, with all four independent runs of the optimization procedure depicted for each body size: (a) mean fitness 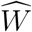, (b) mean maturation time *t*_maturation_, (c-i), the seven different components of the ontogenetic strategy as indicated on each y axis. All solutions are derived with extrinsic mortality *μ* = 0.01, per-division damage risk *c* = 0.01, cell turnover *ν* = 0.0001, number of oncogenic steps *K* = 3, and growth parameter *K* = 0.05.

The remaining four traits (*P, X, S*, and *H*; Fig.2c, f, g, and i, respectively) did not show as clear patterns related to body size. There are two potential explanations: that the fitness landscape is very flat with respect to these traits, or that the traits matter for fitness, but occur in the form of coadapted gene complexes (Sinervo and Svensson, 2002; Schwander et al., 2014). In the latter case, an overall ‘shotgun-like’ pattern (Fig.2c, f, g, and i) has more structure than is apparent at first sight: a rugged multidimensional fitness landscape may produce high fitness at particular combinations of trait values, rather than peaking at a single value for each trait independently of the six other traits. For example, the lower values of *Q* in Fig.2d associate with the higher *P* in Fig.2c. When symmetric divisions contribute less to tissue differentiation (a low *Q*), an organism might not suffer if it resorts to more asymmetric divisions (a high P); at lower *Q*, *P* will matter more (Fig. S1). We tested for the ruggedness of the landscape by using the outcomes of optimizing the ontogenetic strategy four times independently (from different starting points). In case of coevolved gene complexes, creating a new strategy by changing one of the seven evolved components to the outcome of a parallel optimization (for the same body size) should reduce fitness, which we found to be the case (Supplementary material).

We next ask if evolved ontogenetic strategies form constraints for evolving a different body size: can a strategy that evolved at a small body size reach high fitness if implemented in a life history with a larger maturation size; and is the reverse true (from large to small body sizes)? Smaller organisms might fail to operate for long enough if growth to become large (which takes time) is now required. Reciprocally, strategies evolved in larger organisms might be expected to fail if their slow way of forming tissues (low *Q*, and many levels to full differentiation, *T*) means that they are not ‘ready’ when maturing at small size. We found clear evidence for the first type of failure (from small to large) but not the second (Fig.3); strategies evolved in larger organisms were relatively robust to changes in body size, allowing organisms of the entire tested size range to grow to maturation. Interestingly, the independent parallel optimizations for the ‘evolved while small’ strategies varied in the range of size increases that they tolerated (for particularly clear cases see Fig.3b). Thus, while performing equivalently when small, some coadapted combinations permit size increases much better than others.

**Figure 3:**
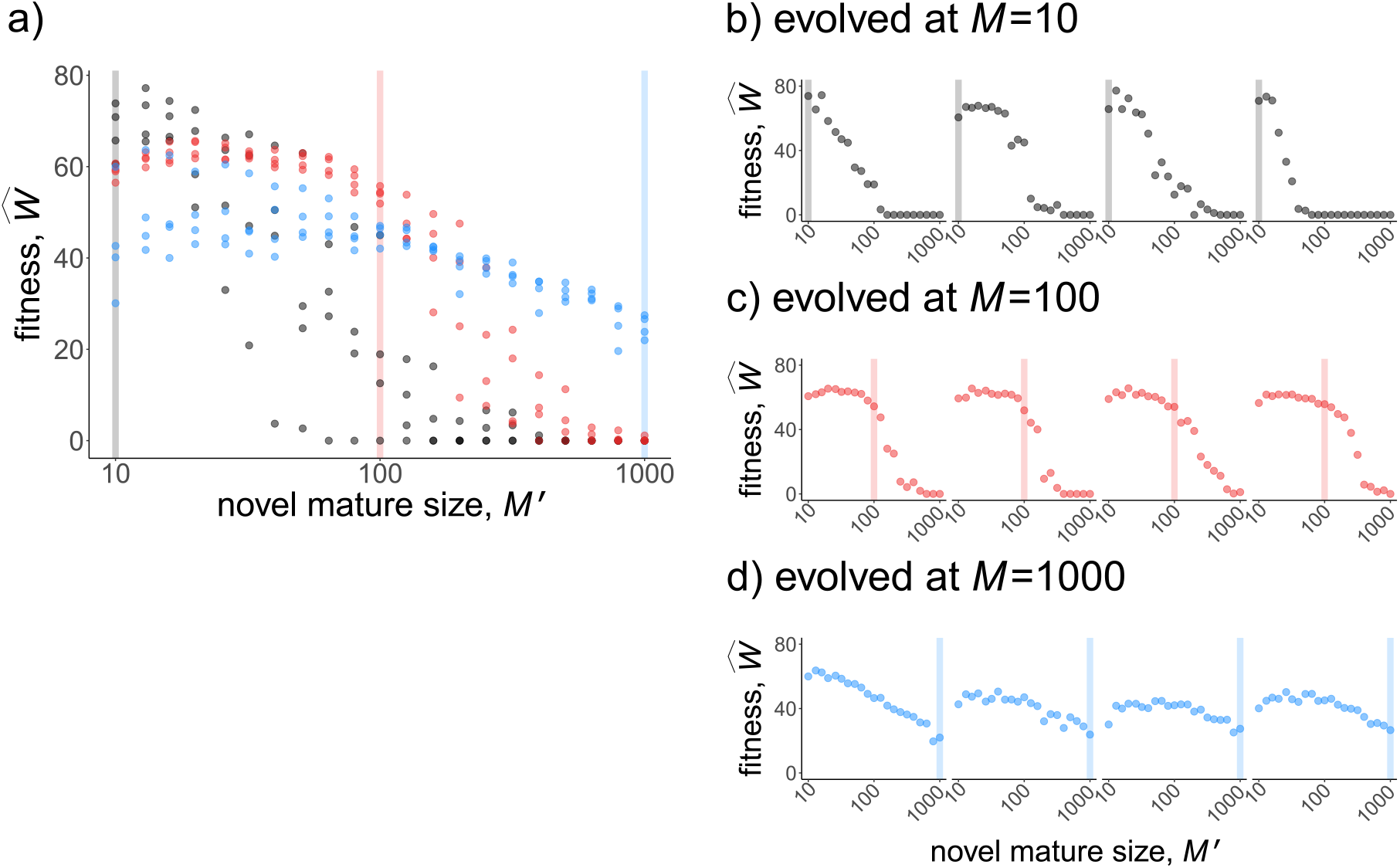
Testing for the performance of ontogenetic strategies evolved at a specific body size at maturity, *M* (vertical lines), at 21 logarithmically spaced novel body sizes (novel maturation size *M*′). In (a), all data is depicted together, to show that strategies evolved under small sizes outperform the others in the range of small sizes (grey), likewise for medium (red) and large (blue) strategies near their own ancestral sizes. In (b, c, d), the same data are divided into the four independent runs, yielding four performance curves that vary considerably in the range of values they can operate in (positive fitness) if tested in organisms differing from the ancestral size (vertical line). Fitness maintenance is easy if the novel body size is small (to the left of the vertical lines in c,d) but hard if large (to the right of the vertical lines in b,c), with the precise shape of the latter pattern being strategy-dependent. All parameters not given in the figure are as in Fig.2.

The causes of death were different across the examined body sizes. Optimized strategies (Fig.4a) typically manage the soma well enough so that no other cause of death than extrinsic mortality occurs until very few individuals are alive (due to extrinsic mortality). In the rare cases where an optimized strategy dies from other causes, then the cause relates to running out of stem cells at small body sizes, and to cancer at large body sizes. However, if a small organism attempts to grow larger (than during its evolutionary past that optimized its ontogenetic strategy), the failures typically relate to running out of stem cells (Fig.4b,c), supporting the interpretation that small organisms do not need as sophisticated management of the stem cell pool as large ones. Large organisms that turn small perform well because the now diminished number of cell divisions protect against cancer, and simultaneously their stem cells do not run out as they were already well managed; extrinsic mortality then remains the virtually only cause of death (left part of Fig.4d).

**Figure 4:**
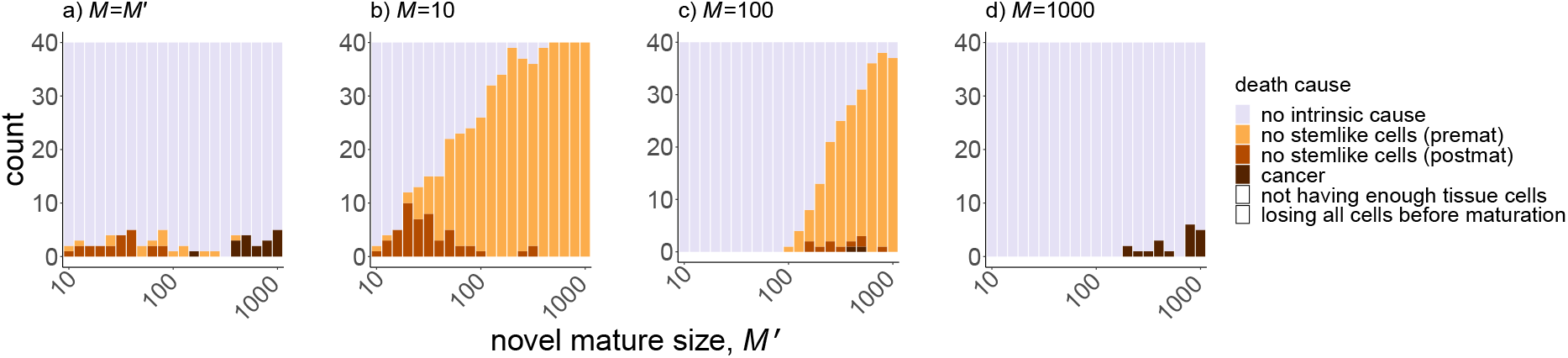
Causes of death (failures of the soma) in different scenarios: (a) when the organism is tested for its own body size (‘novel’ body size equals the one for which the ontogenetic strategy is optimized), (b,c,d) ontogenetic strategy optimized for small, medium and large bodies, respectively, tested across various body sizes. Each scenario is run 40 times (i.e. 10 times for four independently optimized ontogenetic strategies) with the cause of death reported; ‘no intrinsic cause’ implies that the soma was intact for the duration of the entire lifespan when it is assumed to end when extrinsic mortality no longer permits significant survival chances (thus, individuals in this category are predicted to die due to extrinsic causes). In (a) and (d), failures of the soma were rare, while attempts to use an ontogenetic strategy in a larger body (b,c) resulted in a relatively systematic failure to replenish the stem cell population in a sustainable manner. If large body sizes were instead allowed to use a strategy evolved at large size (a,d), cancer replaces stem cell depletion as a probabilistically occurring cause of death. All parameter values are as in figures 2 and 3, unless otherwise indicated. Note that some causes of death (white) never occurred in this test as it used strategies that already had been optimized; they may still have occurred at intermediate steps of the optimization procedure.

### Discussion

Our model integrates seven different aspects of ontogeny (tissue differentiation in the presence of cancer risk) into a single model, to ask (i) how organisms are expected to manage their stem (and stemlike) cell populations during tissue differentiation while also avoiding cancer, and (ii) whether the evolved strategies need to be modified in case organisms change their body size over evolutionary time.

Regarding our first question, it appears that some aspects of division and differentiation strategies are under stronger selection than others, with the fitness landscape as a whole showing signs of ruggedness: for a given body size, different initial trait combinations led to different solutions to the same problem while reaching overall similar fitness. This suggests that organisms of similar size and lifespan, and hence with comparable cell-management requirements, can grow and survive using a diversity of mechanisms. This could potentially explain the variation in telomere length or telomerase expression in mammals, especially in smaller ones (Gomes et al., 2011). While not explicitly modelled by us, it is intriguing that naked mole rats and blind mole rats counteract cancer risk using different mechanisms (Seluanov et al., 2018). Since cancer was only one of several ways to die in our model, our results also highlight the potential for similar ruggedness in stem cell pool replenishment strategies: there are many ways to succeed in keeping the stemlike cell population large enough for long enough, such that the remaining challenges relate to extrinsic mortality only (we here follow a modelling tradition of calling age-independent mortalities that the organism cannot change as ‘extrinsic’; Abrams, 1993; Carnes and Olshansky, 1997; Boddy et al., 2015).

Turning to our second question, the body size dependence of the ontogenetic strategy, we found clear evidence that the strategy requires modification when mature body size changes. However, this finding comes with two subtleties. Firstly, not all components of the strategy showed strong relationships with body size. However, the coadaptedness of strategy components has consequences for the strategies’ robustness when tested in the context of a novel body size. Specifically, forcing two (or more) independently derived strategies to perform at a novel (larger) body size revealed differences in the range of body sizes that a strategy can still cope with. When a strategy failed when tested in a larger body, the problems related to running out of stem cells rather than cancer. This is interesting in the context of Peto’s paradox (Peto et al., 1975; Caulin and Maley, 2011), which is expressible as ‘all else being equal, more cell divisions should lead to a higher cancer risk’. In our model, small organisms died due to stem cell depletion if they became large without being given an opportunity to adapt their ontogeny to their new life history. The cancer risk reappeared only when they used ‘large’-adapted ontogenetic strategies.

Secondly, there was an interesting asymmetry: Large organisms, which adapted to their larger size by having a smaller per-cell probability of differentiation, a not too eagerly applied threshold of killing damaged cells, and many differentiation steps before a tissue is formed, possessed ontogenetic strategies that also worked well when tested in small organisms. Although computational limitations prevented us from scaling our study up to cell numbers involved in large metazoans, we see no reason why such asymmetries would not continue playing a role at such sizes. Our finding is therefore interesting in the context of macroevolutionary changes in body sizes, where it is known that miniaturization, when it happens, appears to proceed much faster than the (as a whole more common; Kingsolver and Pfennig, 2004) increase in body size over evolutionary time (Evans et al., 2012). It also fits well with findings of intraspecific variation in humans, where size increases associate with heightened cancer risk (Green et al., 2011; Nunney, 2018) while hereditary forms of dwarfism, at least in the context of the so-called Laron syndrome, appear to offer cancer protection (Laron et al., 2017).

Note that our model should be complemented with additional considerations when making predictions about rates of evolution. Eco-evolutionary dynamics has features that we did not take into account: larger organisms tend to have smaller effective population sizes (with population density as a proxy; Damuth, 1981), which can slow down the rate of adaptive evolution (Lanfear et al., 2014). We sidestepped such issues as we simply re-simulated optimized ontogenies for novel body sizes, rather than taking existing ontogenies as starting points (while beyond the scope of the current paper, we will examine the consequent phylogenetic inertia in our future work). Generation times also vary with body size (Martin and Palumbi, 1993); our model is an ‘all else being equal’ investigation of body size, where time to maturity takes longer for larger maturation sizes, but e.g. extrinsic mortality does not change. It is interesting to note that our choice to have very few differences between ancestral and novel conditions (no ecological change apart from time it takes to mature, which emerges naturally in the model) still led to a prediction that parallel runs of the optimization procedure will differ in how easily a small-bodied organism can evolve a larger size. Some ontogenetic strategies collapsed to zero fitness at much smaller demands to become big, than others. We interpret this finding as a prediction that some fortuitous small multicellular organisms have preadaptations to grow over macroevolutionary time, while others face stronger intrinsic constraints. This adds ontogenetic challenges to the reasons why Cope’s rule, the increase of body sizes in macroevolutionary lineages, is expected to be a rather stochastic ‘rule’ (see e.g. Liow and Taylor, 2019).

Some of our findings are, at least at first sight, unexpected. The damage response threshold *A* reflects how many mutations are tolerated before a cell is recognized as damaged and killed (e.g. via apoptosis, or by other cells; we do not specify the exact mechanism) with probability *S*. Based on an important study on elephant cells responding more readily to damage than human cells (Abegglen et al., 2015), one might expect *A* to decrease and *S* to increase with body size, but such responses did not emerge in our model. Instead, small organisms could function across a wide range of values for both *A* and *S*, while larger somas were never found to be built with small values of *A*. This apparent mismatch between Abegglen et al. (2015) and our results disappears, however, when taking into account coadaptation between different ontogenetic strategy components. Large organisms also evolved to use more steps (*T*) to full differentiation, which also gives cell lineages more cumulative opportunities (in the form of a higher number of divisions) to accumulate damage. A higher *A*, when it also associates with a higher *T*, therefore does not necessarily mean that fewer cells are observed to be killed in any phenotype. When *T* is high, cells have more opportunities to reach a category of damage that sees them killed. Moreover, since running out of stemlike cells was a real threat in our model, it appears that this penalizes against very strict ‘quality control’ of cells; no strategy can succeed if sufficient cell numbers are not maintained, while cancer can at least probabilistically be avoided (under a multi-step model of oncogenesis; Armitage and Doll, 1954; Nunney, 1999; Calabrese and Shibata, 2010). These considerations add to our general message that there are many complementary ways to create a soma of a certain body size.

## Acknowledgements

This work was supported by the University Research Priority Program (URPP) “Evolution in Action” of the University of Zurich. Authors thank Arizona Cancer Evolution Centre (ACE) members for insights.

## Data availability statement

Simulation code used in this study is publicly available in GitHub: https://github.com/yagmurerten/OptimiseOntogeny.

## References

Abegglen, L. M., Caulin, A. F., Chan, A., Lee, K., Robinson, R., Campbell, M. S., Kiso, W. K., Schmitt, D. L., Waddell, P. J., Bhaskara, S., Jensen, S. T., Maley, C. C. and Schiffman, J. D. (2015). Potential Mechanisms for Cancer Resistance in Elephants and Comparative Cellular Response to DNA Damage in Humans, JAMA 314(17): 1850–1860. doi:10.1001/jama.2015.13134

Abrams, P. A. (1993). Does increased mortality favor the evolution of more rapid senescence?, Evolution 47(3): 877–887. doi:10.1111/j.1558-5646.1993.tb01241.x

Aktipis, C. A., Boddy, A. M., Jansen, G., Hibner, U., Hochberg, M. E., Maley, C. C. and Wilkinson, G. S. (2015). Cancer across the tree of life: cooperation and cheating in multicellularity, Philosophical Transactions of the Royal Society B: Biological Sciences 370(1673): 20140219. doi:10.1098/rstb.2014.0219

Armitage, P. and Doll, R. (1954). The age distribution of cancer and a multi-stage theory of carcinogenesis., British Journal of Cancer 8(1): 1–12. doi:10.1038/bjc.1954.1pmid:13172380

Bastiaans, E., Debets, A. J. M. and Aanen, D. K. (2016). Experimental evolution reveals that high relatedness protects multicellular cooperation from cheaters, Nature Communications 7(1): 11435. doi:10.1038/ncomms11435

Blasco, M. A. (2005). Telomeres and human disease: ageing, cancer and beyond, Nature Reviews Genetics 6(8): 611–622. doi:10.1038/nrg1656

Boddy, A. M., Kokko, H., Breden, F., Wilkinson, G. S. and Aktipis, C. A. (2015). Cancer susceptibility and reproductive trade-offs: a model of the evolution of cancer defences, Philosophical Transactions of the Royal Society B: Biological Sciences 370(1673): 20140220. doi:10.1098/rstb.2014.0220

Boraas, M. E., Seale, D. B. and Boxhorn, J. E. (1998). Phagotrophy by a flagellate selects for colonial prey: A possible origin of multicellularity, Evolutionary Ecology 12(2): 153–164. doi:10.1023/A:1006527528063

Calabrese, P. and Shibata, D. (2010). A simple algebraic cancer equation: calculating how cancers may arise with normal mutation rates, BMC Cancer 10(1): 3. doi:10.1186/1471-2407-10-3

Campisi, J. (2013). Aging, cellular senescence, and cancer, Annual Review of Physiology 75(1): 685–705. PMID: 23140366. doi:10.1146/annurev-physiol-030212-183653

Carnes, B. A. and Olshansky, S. (1997). A biologically motivated partitioning of mortality, Experimental Gerontology 32(6): 615–631. doi:10.1016/S0531-5565(97)00056-9

Caulin, A. F. and Maley, C. C. (2011). Peto’s Paradox: evolution’s prescription for cancer prevention, Trends in Ecology & Evolution 26(4): 175–182. doi:10.1016/j.tree.2011.01.002

Damuth, J. (1981). Population density and body size in mammals, Nature 290(5808): 699–700. doi:10.1038/290699a0

Derényi, I. and Szöllősi, G. J. (2017). Hierarchical tissue organization as a general mechanism to limit the accumulation of somatic mutations, Nature Communications 8(1): 14545. doi:10.1038/ncomms14545

Evans, A. R., Jones, D., Boyer, A. G., Brown, J. H., Costa, D. P., Ernest, S. K. M., Fitzgerald, E. M. G., Fortelius, M., Gittleman, J. L., Hamilton, M. J., Harding, L. E., Lintulaakso, K., Lyons, S. K., Okie, J. G., Saarinen, J. J., Sibly, R. M., Smith, F. A., Stephens, P. R., Theodor, J. M. and Uhen, M. D. (2012). The maximum rate of mammal evolution, Proceedings of the National Academy of Sciences 109(11): 4187–4190. doi:10.1073/pnas.1120774109

Fisher, R., Cornwallis, C. and West, S. (2013). Group Formation, Relatedness, and the Evolution of Multicellularity, Current Biology 23(12): 1120–1125. doi:10.1016/j.cub.2013.05.004

Fuchs, Y. and Steller, H. (2011). Programmed cell death in animal development and disease, Cell 147(4): 742–758. doi:https://doi.org/10.1016/j.cell.2011.10.033

Galluzzi, L., Vitale, I., Aaronson, S. A., Abrams, J. M., Adam, D., Agostinis, P., Alnemri, E. S., Altucci, L., Amelio, I., Andrews, D. W., Annicchiarico-Petruzzelli, M., Antonov, A. V., Arama, E., Baehrecke, E. H., Barlev, N. A., Bazan, N. G., Bernassola, F., Bertrand, M. J. M., Bianchi, K., Blagosklonny, M. V., Blomgren, K., Borner, C., Boya, P., Brenner, C., Campanella, M., Candi, E., Carmona-Gutierrez, D., Cecconi, F., Chan, F. K.-M., Chandel, N. S., Cheng, E. H., Chipuk, J. E., Cidlowski, J. A., Ciechanover, A., Cohen, G. M., Conrad, M., Cubillos-Ruiz, J. R., Czabotar, P. E., D’Angiolella, V., Dawson, T. M., Dawson, V. L., De Laurenzi, V., De Maria, R., Debatin, K.-M., DeBerardinis, R. J., Deshmukh, M., Di Daniele, N., Di Virgilio, F., Dixit, V. M., Dixon, S. J., Duckett, C. S., Dynlacht, B. D., El-Deiry, W. S., Elrod, J. W., Fimia, G. M., Fulda, S., García-Saez, A. J., Garg, A. D., Garrido, C., Gavathiotis, E., Golstein, P., Gottlieb, E., Green, D. R., Greene, L. A., Gronemeyer, H., Gross, A., Hajnoczky, G., Hardwick, J. M., Harris, I. S., Hengartner, M. O., Hetz, C., Ichijo, H., Jäättela, M., Joseph, B., Jost, P. J., Juin, P. P., Kaiser, W. J., Karin, M., Kaufmann, T., Kepp, O., Kimchi, A., Kitsis, R. N., Klionsky, D. J., Knight, R. A., Kumar, S., Lee, S. W., Lemasters, J. J., Levine, B., Linkermann, A., Lipton, S. A., Lockshin, R. A., Líopez-Otín, C., Lowe, S. W., Luedde, T., Lugli, E., MacFarlane, M., Madeo, F., Malewicz, M., Malorni, W., Manic, G., Marine, J.-C., Martin, S. J., Martinou, J.-C., Medema, J. P., Mehlen, P., Meier, P., Melino, S., Miao, E. A., Molkentin, J. D., Moll, U. M., Muñoz-Pinedo, C., Nagata, S., Nuñez, G., Oberst, A., Oren, M., Overholtzer, M., Pagano, M., Panaretakis, T., Pasparakis, M., Penninger, J. M., Pereira, D. M., Pervaiz, S., Peter, M. E., Piacentini, M., Pinton, P., Prehn, J. H., Puthalakath, H., Rabinovich, G. A., Rehm, M., Rizzuto, R., Rodrigues, C. M., Rubinsztein, D. C., Rudel, T., Ryan, K. M., Sayan, E., Scorrano, L., Shao, F., Shi, Y., Silke, J., Simon, H.-U., Sistigu, A., Stockwell, B. R., Strasser, A., Szabadkai, G., Tait, S. W., Tang, D., Tavernarakis, N., Thorburn, A., Tsujimoto, Y., Turk, B., Vanden Berghe, T., Vandenabeele, P., Vander Heiden, M. G., Villunger, A., Virgin, H. W., Vousden, K. H., Vucic, D., Wagner, E. F., Walczak, H., Wallach, D., Wang, Y., Wells, J. A., Wood, W., Yuan, J., Zakeri, Z., Zhivotovsky, B., Zitvogel, L., Melino, G. and Kroemer, G. (2018). Molecular mechanisms of cell death: recommendations of the Nomenclature Committee on Cell Death 2018, Cell Death & Differentiation 25(3): 486–541. doi:10.1038/s41418-017-0012-4

Goldsby, H. J., Knoester, D. B., Ofria, C. and Kerr, B. (2014). The Evolutionary Origin of Somatic Cells under the Dirty Work Hypothesis, PLoS Biology 12(5): e1001858. doi:10.1371/journal.pbio.1001858

Gomes, N. M. V., Ryder, O. A., Houck, M. L., Charter, S. J., Walker, W., Forsyth, N. R., Austad, S. N., Venditti, C., Pagel, M., Shay, J. W. and Wright, W. E. (2011). Comparative biology of mammalian telomeres: hypotheses on ancestral states and the roles of telomeres in longevity determination: The comparative biology of mammalian telomeres, Aging Cell 10(5): 761–768. doi:10.1111/j.1474-9726.2011.00718.x

Green, J., Cairns, B. J., Casabonne, D., Wright, F. L., Reeves, G. and Beral, V. (2011). Height and cancer incidence in the Million Women Study: prospective cohort, and meta-analysis of prospective studies of height and total cancer risk, The Lancet Oncology 12(8): 785–794. doi:10.1016/S1470-2045(11)70154-1

Grosberg, R. K. and Strathmann, R. R. (2007). The Evolution of Multicellularity: A Minor Major Transition?, Annual Review of Ecology, Evolution, and Systematics 38(1): 621–654. doi:10.1146/annurev.ecolsys.36.102403.114735

Hayflick, L. (1965). The limited in vitro lifetime of human diploid cell strains, Experimental Cell Research 37(3): 614–636. doi:https://doi.org/10.1016/0014-4827(65)90211-9

Hayflick, L. and Moorhead, P. (1961). The serial cultivation of human diploid cell strains, Experimental Cell Research 25(3): 585–621. doi:10.1016/0014-4827(61)90192-6

Kapsetaki, S. E. and West, S. A. (2019). The costs and benefits of multicellular group formation in algae*, Evolution 73(6): 1296–1308. doi:10.1111/evo.13712

Karkach, A. S. (2006). Trajectories and models of individual growth, Demographic Research 15: 347–400. doi:10.4054/DemRes.2006.15.12

Kim, N., Piatyszek, M., Prowse, K., Harley, C., West, M., Ho, P., Coviello, G., Wright, W., Weinrich, S. and Shay, J. (1994). Specific association of human telomerase activity with immortal cells and cancer, Science 266(5193): 2011–2015. doi:10.1126/science.7605428

Kingsolver, J. G. and Pfennig, D. W. (2004). Individual-level selection as a cause of Cope’s rule of phyletic size increase, Evolution 58(7): 1608–1612. doi:10.1111/j.0014-3820.2004.tb01740.x

Knoll, A. H. (2011). The Multiple Origins of Complex Multicellularity, Annual Review of Earth and Planetary Sciences 39(1): 217–239. doi:10.1146/annurev.earth.031208.100209

Kokko, H. and Hochberg, M. E. (2015). Towards cancer-aware life-history modelling, Philosophical Transactions of the Royal Society B: Biological Sciences 370(1673): 20140234. doi:10.1098/rstb.2014.0234

Lanfear, R., Kokko, H. and Eyre-Walker, A. (2014). Population size and the rate of evolution, Trends in Ecology & Evolution 29(1): 33–41. doi:10.1016/j.tree.2013.09.009

Laron, Z., Kauli, R., Lapkina, L. and Werner, H. (2017). IGF-I deficiency, longevity and cancer protection of patients with Laron syndrome, Mutation Research/Reviews in Mutation Research 772: 123–133. doi:https://doi.org/10.1016/j.mrrev.2016.08.002

Libby, E., Conlin, P. L., Kerr, B. and Ratcliff, W. C. (2016). Stabilizing multicellularity through ratcheting, Philosophical Transactions of the Royal Society B: Biological Sciences 371(1701): 20150444. doi:10.1098/rstb.2015.0444

Liow, L. H. and Taylor, P. D. (2019). Cope’s rule in a modular organism: Directional evolution without an overarching macroevolutionary trend, Evolution 73(9): 1863–1872. doi:10.1111/evo.13800

Liu, N., Matsumura, H., Kato, T., Ichinose, S., Takada, A., Namiki, T., Asakawa, K., Morinaga, H., Mohri, Y., De Arcangelis, A., Geroges-Labouesse, E., Nanba, D. and Nishimura, E. K. (2019). Stem cell competition orchestrates skin homeostasis and ageing, Nature 568(7752): 344–350. doi:10.1038/s41586-019-1085-7

Maciejowski, J. and de Lange, T. (2017). Telomeres in cancer: tumour suppression and genome instability, Nature Reviews Molecular Cell Biology 18(3): 175–186. doi:10.1038/nrm.2016.171

Martin, A. P. and Palumbi, S. R. (1993). Body size, metabolic rate, generation time, and the molecular clock., Proceedings of the National Academy of Sciences 90(9): 4087–4091. doi:10.1073/pnas.90.9.4087

Nunney, L. (1999). Lineage selection and the evolution of multistage carcinogenesis, Proceedings of the Royal Society of London. Series B: Biological Sciences 266(1418): 493–498. doi:10.1098/rspb.1999.0664

Nunney, L. (2018). Size matters: height, cell number and a person’s risk of cancer, Proceedings of the Royal Society B: Biological Sciences 285(1889): 20181743. doi:10.1098/rspb.2018.1743

Peto, R., Roe, F. J., Lee, P. N., Levy, L. and Clack, J. (1975). Cancer and ageing in mice and men, British Journal of Cancer 32(4): 411–426. doi:10.1038/bjc.1975.242

Pettersen, A. K., White, C. R. and Marshall, D. J. (2015). Why does offspring size affect performance? Integrating metabolic scaling with life-history theory, Proceedings of the Royal Society B: Biological Sciences 282(1819): 20151946. doi:10.1098/rspb.2015.1946

Queller, D. C. (2000). Relatedness and the fraternal major transitions, Philosophical Transactions of the Royal Society of London. Series B: Biological Sciences 355(1403): 1647–1655. doi:10.1098/rstb.2000.0727

Risques, R. A. and Promislow, D. E. L. (2018). All’s well that ends well: why large species have short telomeres, Philosophical Transactions of the Royal Society B: Biological Sciences 373(1741): 20160448. doi:10.1098/rstb.2016.0448

Sager, R. (1991). Senescence as a mode of tumor suppression, Environmental Health Perspectives 93: 59–62. doi:10.1289/ehp.919359

Schwander, T., Libbrecht, R. and Keller, L. (2014). Supergenes and Complex Phenotypes, Current Biology 24(7): R288–R294. doi:https://doi.org/10.1016/j.cub.2014.01.056

Seluanov, A., Chen, Z., Hine, C., Sasahara, T. H. C., Ribeiro, A. A. C. M., Catania, K. C., Presgraves, D. C. and Gorbunova, V. (2007). Telomerase activity coevolves with body mass not lifespan, Aging Cell 6(1): 45–52. doi:10.1111/j.1474-9726.2006.00262.x

Seluanov, A., Gladyshev, V. N., Vijg, J. and Gorbunova, V. (2018). Mechanisms of cancer resistance in long-lived mammals, Nature Reviews Cancer 18(7): 433–441. doi:10.1038/s41568-018-0004-9

Shahriyari, L. and Komarova, N. L. (2013). Symmetric vs. Asymmetric Stem Cell Divisions: An Adaptation against Cancer?, PLOS ONE 8(10): 1–16. doi:10.1371/journal.pone.0076195

Shay, J. W. and Wright, W. E. (2019). Telomeres and telomerase: three decades of progress, Nature Reviews Genetics 20(5): 299–309. doi:10.1038/s41576-019-0099-1

Sinervo, B. and Svensson, E. (2002). Correlational selection and the evolution of genomic architecture, Heredity 89(5): 329–228. doi:10.1038/sj.hdy.6800148

White, C. R., Marshall, D. J., Alton, L. A., Arnold, P. A., Beaman, J. E., Bywater, C. L., Condon, C., Crispin, T. S., Janetzki, A., Pirtle, E., Winwood-Smith, H. S., Angilletta, M. J., Chenoweth, S. F., Franklin, C. E., Halsey, L. G., Kearney, M. R., Portugal, S. J. and Ortiz-Barrientos, D. (2019). The origin and maintenance of metabolic allometry in animals, Nature Ecology & Evolution 3(4): 598–603. doi:10.1038/s41559-019-0839-9

