## Supporting Information for "From zygote to a multicellular soma: body size affects optimal growth strategies under cancer risk"

### Testing for coadaptation of traits within strategies

In the main text, we reported that only a few traits (namely, probability of differentiation $Q$, number of differentiation steps $T$, and damage response threshold $A$) evolved clear systematic differences across body sizes, whereas the other four (probability of asymmetric divisions $P$, Hayflick limit $H$, division propensity $X$, and damage response strength $S$) could vary widely while yielding equivalently high fitness values. While one hypothesis is that the latter four traits relate only weakly to the fitness of an ontogenic management strategy (thus evolving largely neutrally in a genetic algorithm), the alternative is that some or all seven traits could be coadapted to each other with different combinations leading to similar fitness through complementation (e.g. Fig.S1). In the former case, the fitness of already optimized solutions should not change in any systematic manner if one of the seven components from one strategy combines with other components being chosen from another optimized strategy (both optimizations having been perfomed under the same conditions such as body size, but from different starting conditions). Taking one component from a competing solution would be expected to yield either higher or lower fitness, depending on which value is closer to the global optimum. However, this expectation changes if there are coadapted gene complexes. In that case, changing the value of one component to one from a competing solution is expected to lead to lower fitness.

To test this, we created a set of 'disrupted strategies' by switching the value of one trait with the value (for the same trait) evolved as part of another strategy optimised for the same body size. As we had four competing solutions for each body size, and each strategy has three competitors from where an alternative trait value can be chosen to disrupt a potentially coadapted gene complex, the trials involved 12 disrupted strategies per trait for each body size. For each trait and body size combination, we also estimated fitness of the optimized strategies themselves; as this involves not changing any traits, we essentially re-estimate the fitness of the optimal strategies. For both the disrupted and control strategies, we ran 10 replicates to estimate the mean fitness ($\widehat{W}_1$ and $\widehat{W}_0$, respectively). We considered a negative $r = \ln(\widehat{W}_1/\widehat{W}_0)$ as evidence for coadapted gene complexes.

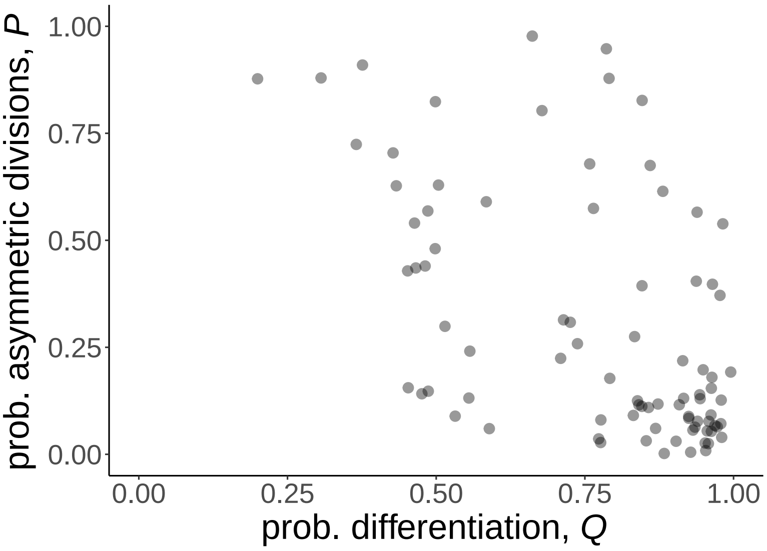


Figure S1: Relationship between probabilities of asymmetric division ($P$) and differentiation ($Q$). A low probability of differentiation associates with a high probability of asymmetric divisions. Parameters and the body sizes simulated are as in Fig.2 of the main text.

Our results showed that in approximately 3% of the cases, changing a single trait within an optimised strategy rendered it completely unsuccessful, leading to zero fitness (50 cases out of 1764, with $r = -\infty$, Fig.S2). Only two traits, the Hayflick limit $H$ and level-dependence of the division propensity $X$, avoided having any of these disrupted cases; thus, all other traits showed evidence of the trait playing a part of coadaptation with other components of the ontogenetic strategy.

We thereafter proceeded to analyze the remaining 97% of cases where the difference effect of disruption was milder (i.e. $\widehat{W}_1$ remained positive). Clear evidence for a disrupted strategy performing on average worse, when conservatively having first removed the complete failures, was found for the probability of asymmetric divisions $P$ (paired one-sided t-test, t=−6.508, df=246, p=$2.117\times10^{-10}$), probability of differentiation $Q$ (t=−6.806, df=244, p=$3.870\times10^{-11}$), the number of differentiation steps $T$ (t=−5.367, df=215, p=$1.033\times10^{-7}$) and the relative division propensity of differentiation levels $X$ (t=−3.124, df=251, p=0.001); note that these remain significant after a Bonferroni correction (which for 7 comparisons demands an $\alpha$ value of 0.0071 or lower).

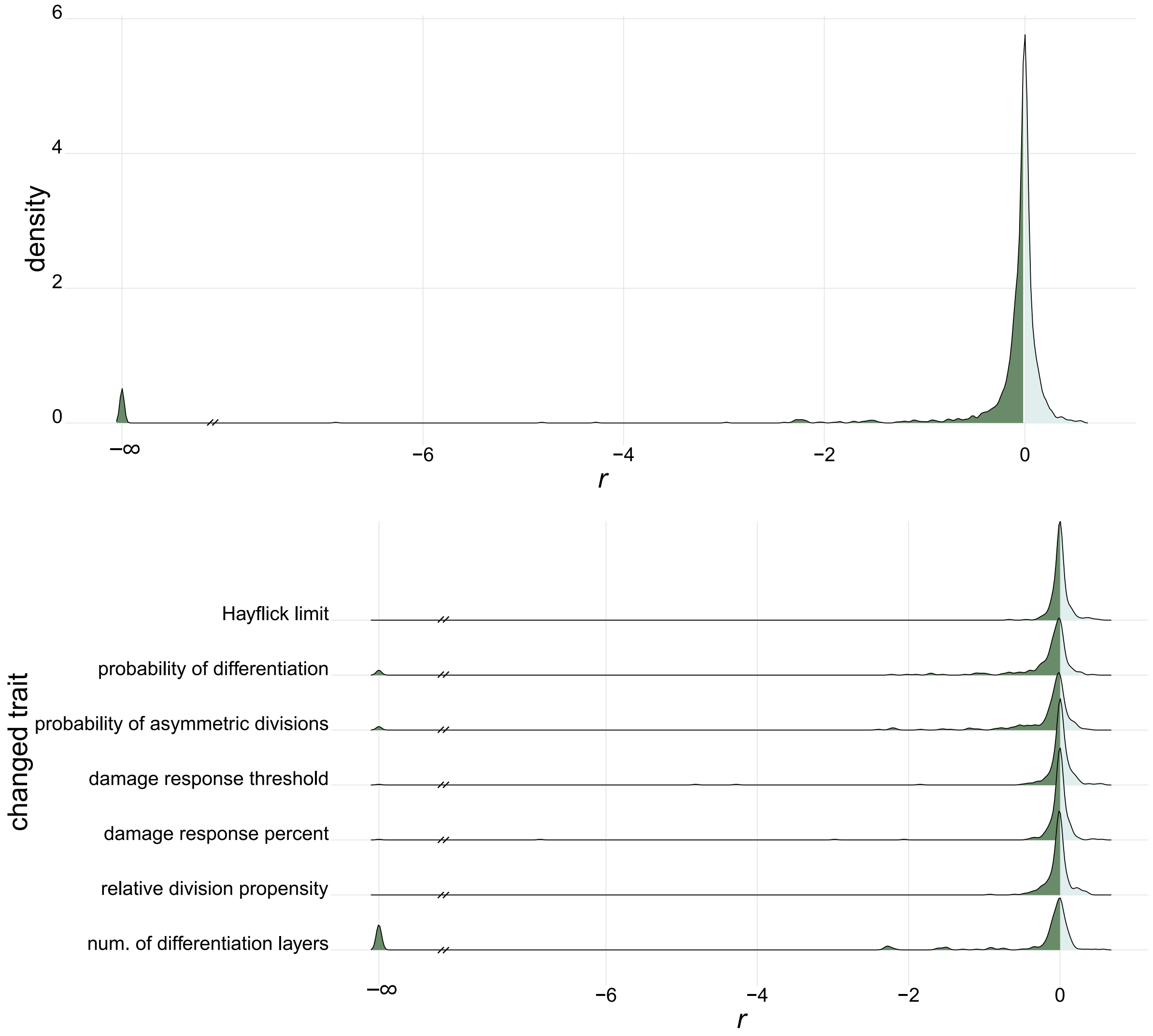


Figure S2: Performance of disrupted strategies. While it is not impossible that a disrupted strategy outperforms the original (lighter shaded cases: positive $r$), on average there are more cases where the fitness declines (darker shaded cases: negative $r$). The upper panel shows all data combined, and the lower panel subdivides the data into trait-specific categories.
